# Mitochondrial morphology dynamics and ROS regulate apical polarity and differentiation in *Drosophila* follicle cells

**DOI:** 10.1101/2023.03.10.532033

**Authors:** Bhavin Uttekar, Darshika Tomer, Richa Rikhy

## Abstract

Mitochondrial morphology dynamics regulate signaling pathways during epithelial cell formation and differentiation. The mitochondrial fission protein Drp1 affects the appropriate activation of EGFR and Notch signaling-driven differentiation of posterior follicle cells in *Drosophila* oogenesis. The mechanisms by which Drp1 regulates epithelial polarity during differentiation are not known. In this study, we show that Drp1 depleted follicle cells are constricted in early stages and present in multiple layers at later stages with decreased levels of apical polarity protein aPKC. This defect is suppressed by additional depletion of mitochondrial fusion protein Opa1. Opa1 depletion leads to mitochondrial fragmentation and increased reactive oxygen species (ROS) in follicle cells. We find that increasing ROS by depleting the ROS scavengers, mitochondrial SOD2, and catalase also leads to mitochondrial fragmentation. Further, the loss of Opa1, SOD2, and catalase partially restores the defects in epithelial polarity and aPKC along with EGFR and Notch signaling in Drp1 depleted follicle cells. Our results show a crucial interaction between mitochondrial morphology, ROS generation, and epithelial cell polarity formation during the differentiation of follicle epithelial cells in *Drosophila* oogenesis.

**Summary statement:** Mitochondrial fission protein Drp1 regulates epithelial follicle cell differentiation in *Drosophila* oogenesis. Increasing ROS and mitochondrial fragmentation suppresses the defects in epithelial polarity, and differentiation in Drp1 depleted follicle cells.

## Introduction

Signaling pathways regulate the onset and remodeling of epithelial cell polarity in stem cell differentiation and embryogenesis. Epithelial cells have a polarized plasma membrane organization with an apical, lateral, and basal domain containing different protein complexes. Epithelial cell differentiation accompanies the presence of elaborate and fused mitochondria in mammalian hepatocytes. A critical role for mitochondrial fusion-derived OXPHOS-dependent energy has been seen in epithelial polarity establishment (Fu et al., 2013). Mammalian MDCK cells need an ATP-rich environment for the integrity of junctional complexes (Tsukamoto and Nigam, 1997; Tsukamoto and Nigam, 1999). These studies show a role for mitochondrial morphology and associated metabolites in the form of ATP in regulating polarity formation and maintenance (Madan et al., 2021). However, the mechanisms by which polarity complexes are regulated by mitochondrial morphology dynamics and function during stem cell differentiation remain to be investigated.

An immature apical domain is present at the stem cell stage compared to the differentiated stage in certain epithelial cells in *Drosophila* and mammalian systems. *Drosophila* intestinal stem cells lack an apical domain. A mature apical domain is formed on the differentiation of these cells into enteroblasts and enterocytes. The inhibition of mitochondrial fusion protein Opa1 (Deng et al., 2018) and mitochondrial respiratory (ETC) complexes reduces intestinal stem cell differentiation into enterocytes (Zhang et al., 2020). Thus, epithelial cell differentiation is influenced by mitochondrial morphology and function.

*Drosophila* follicle stem cell (FSC) is another example of a partially polarized epithelial cell that lacks an apical domain (Castanieto et al., 2014). In this study, we have analyzed the role of regulation of mitochondrial dynamics in apical polarity maintenance in late *Drosophila* follicle cell (FC) differentiation. FCs encase germ cell cysts during *Drosophila* oogenesis. The germline stem cells (GSCs) and FSCs are located in the germarium (Fadiga and Nystul, 2019). Each germline stem cell undergoes 4 incomplete divisions and forms a 16-cell cyst. Each FSC divides and forms pre-follicle cells which further divide to form FCs (Castanieto et al., 2014; Ulmschneider et al., 2016). FSCs lack an apical domain whereas mature follicle epithelial cells possess apical, basolateral, and basal domains characterized by a domain-specific set of polarity proteins (Castanieto et al., 2014; Dobens and Raftery, 2000; Wu et al., 2008). The apical domain has the aPKC-PAR3-PAR6 complex and the basolateral domain has the Dlg-Scribble-Lgl complex. The apical and basolateral domains are separated by the adherens junction complex (Kaplan et al., 2009). FCs undergo mitosis in stages 1-6 and enter the endocycle from stage 6 by activating the Notch signaling pathway (Klusza and Deng, 2011) (Fig. 1A). The mitotic stage FCs express the homeodomain transcription factor Cut (Sun and Deng, 2005). Transitioning from the mitotic stage to the endocycling stage requires the activation of the Notch signaling pathway and the expression of the transcription factor Hindsight (Jia et al., 2015; Klusza and Deng, 2011; Shcherbata et al., 2004). An active EGFR signaling pathway inhibits the formation of the apical domain in FSCs whereas newly formed pre-FCs show an apical domain due to the suppression of EGFR signaling (Castanieto et al., 2014). Activation of the EGFR signaling pathway also regulates differentiation of the posterior follicle cells (PFCs) at late stages. The stepwise polarity establishment in the follicle epithelial cells during *Drosophila* oogenesis is an insightful model for studying the mechanisms that regulate epithelial cell differentiation and polarity establishment (Franz and Riechmann, 2010; Tepass et al., 2001).

**Figure 1:**
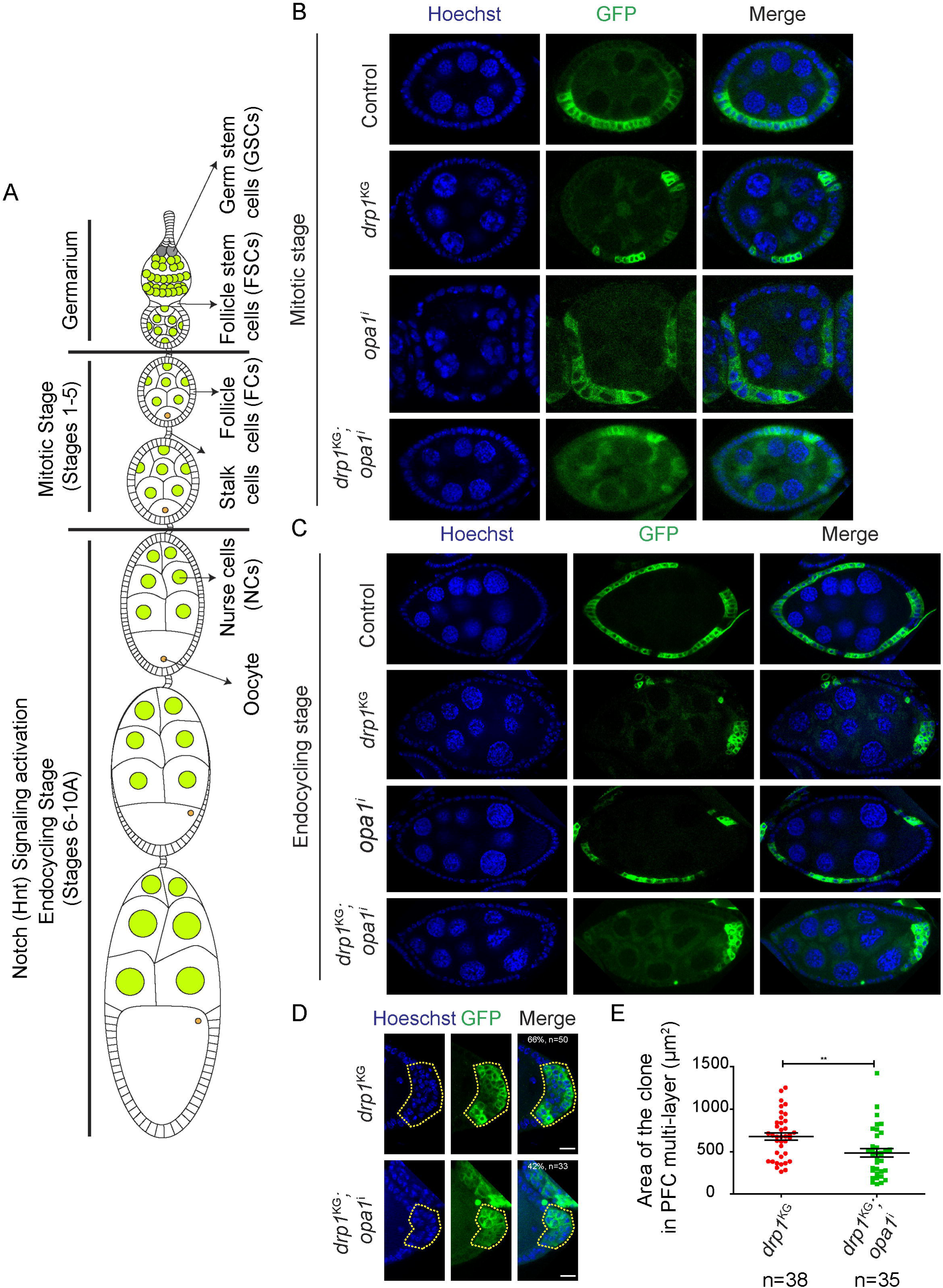
Multilayering in Drp1 depleted PFCs is reduced by additional depletion of Opa1. **(A-E)** Schematic diagram showing *Drosophila* ovariole and stages of egg chambers **(A)**. Representative images of FC arrangement in control FRT40A, *drp1*^KG^, *opa1*^i^, and *drp1*^KG^;*opa1*^i^ follicle cell clones at mitotic stage **(B)**. Representative images of FC arrangement in control FRT40A, *drp1*^KG^, *opa1*^i^, and *drp1*^KG^;*opa1*^i^ PFC clones at endocycling stage **(C)**. Representative images showing multiple layers of PFC clones in *drp1*^KG^ (66% clones have 3 layers or more, n=50) and *drp1*^KG^;*opa1*^i^ (42% clones have 3 layers or more, n=33) (also see Fig. S1C) **(D)**. The graph shows a comparison of the PFC clone area between *drp1*^KG^ (n=38) and *drp1*^KG^;*opa1*^i^ (n=35) **(E)**. The plot shows Mean±SEM. Each data point (n) represents a clone in a separate egg chamber. The statistical test performed for clone area measurement is Student’s t-test. ns=non-significant, * P<0.05, ** P<0.01, *** P<0.001. mCD8-GFP (green) expressing FC clones of the indicated genotype are marked with dashed yellow outlines. The nucleus (blue) is stained with Hoechst. Scale bar=10μm.

FCs show an increase in mitochondrial density during *Drosophila* oogenesis (Tourmente et al., 1990). They show dispersed mitochondria at early stages (Cox and Spradling, 2009). FSCs are lost in mutants causing mitochondrial dysfunction and in mutants leading to an increase in reactive oxygen species (ROS) (Wang et al., 2012). Blocking mitochondrial fission by depleting Drp1 leads to fused mitochondria and this inhibits the transition to the endocycling stage in *Drosophila* oogenesis (Mitra et al., 2012; Tomer et al., 2018). The Drp1 depleted PFCs show an accumulation of phosphorylated ERK downstream of EGFR signaling and loss of Notch signaling. They are present in multiple layers. In this study, we assess the mechanism by which mitochondrial morphology regulates epithelial cell polarity maintenance during *Drosophila* oogenesis. We find that Drp1 deficient FCs show depletion of apical polarity protein aPKC even before the formation of multilayers. This defect is suppressed by the fragmentation of mitochondria by additional depletion of mitochondrial fusion protein Opa1 in Drp1 depleted FCs. An increase in ROS by Opa1 depletion or inhibition of ROS scavenging enzymes leads to mitochondrial fragmentation and the formation of apical polarity in Drp1 depleted FCs. The increase in ROS also leads to suppression of the accumulation of cytoplasmic ERK and loss of Notch-mediated differentiation defect in Drp1 depleted FCs. Our study shows an interaction of mitochondrial dynamics in the regulation of the levels of ROS and follicle epithelial cell formation in *Drosophila* oogenesis.

## Materials and Methods

### Drosophila Genetics

*Drosophila* strains were grown and maintained at 25 °C on a standard cornmeal agar medium. The *Drosophila* stocks obtained from the Bloomington Stock Center are *drp1*^KG^ (y[1]; P{y[+mDint2] w[BR.E.BR]=SUPor-P}Drp1[KG03815]/CyO; ry[506], Bloomington stock number #(BL)13510) FRT40A, UAS-*opa1* RNAi (*opa1*^i^) (y[1] sc[*] v[1] sev[21]; P{y[+t7.7] v[+t1.8]=TRiP.HMS00349}attP2, # BL32358) UAS-*erk* RNAi (*erk*^i^) (y[1] sc[*] v[1] sev[21]; P{y[+t7.7] v[+t1.8]=TRiP.HMS00173}attP2, #BL34855), UAS-*sod2* RNAi (*sod2*^i^) (y[1] v[1]; P{y[+t7.7] v[+t1.8]=TRiP.JF01989}attP2, # BL25969), UAS-*catalase* RNAi (*catalase*^i^) (y[1] sc[*] v[1] sev[21]; P{y[+t7.7] v[+t1.8]=TRiP.HMS00990}attP2, #BL34020). For the MARCM experiments, *hs*-FLP; Gal80-FRT40A/CyO; *tub*-Gal4,UAS-CD8-GFP/TM6, and FRT40A/SM6a were obtained from Nicole Grieder and Mary Lilly, Bethesda, NIH, USA. Standard genetic crosses were performed to make the following combinations for experiments:

*drp1*^KG^ FRT40A/ CyO; *opa1*^i^ 32358/TM6 and FRT40A/ CyO; *opa1*^i^ 32358/TM6,

*drp1*^KG^ FRT40A/ CyO; *erk*^i^ 34855/TM6 and FRT40A/ CyO; *erk*^i^ 34855/TM6,

*drp1*^KG^ FRT40A/ CyO; *sod2*^i^ 25969/TM6 and FRT40A/ CyO; *sod2*^i^ 25969/TM6,

*drp1*^KG^ FRT40A/ CyO; *catalase*^i^ 34020/TM6 and FRT40A/ CyO; *catalase*^i^ 34020/TM6,

### Induction of the mitotic clones using the MARCM technique

The flies having genotype *drp1*^KG^ FRT40A/CyO were crossed to *hs*-FLP; Gal80 FRT40A/CyO; *tub*-Gal4,UAS-CD8-GFP/TM6. F1 flies having the genotype *hs*-FLP/+; *drp1*^KG^ FRT40A/ Gal80 FRT40A; *tub*-Gal4,UAS-CD8-GFP/+ were heat shocked at 37.5 °C for 60 minutes in the water bath to induce follicle cell clones in the ovaries. The flies were transferred to the vials containing fresh media with yeast granules and were further transferred into fresh media every 3 days until they aged 10 days to assess germline clones (i.e 10 days post heat shock). To perform RNAi-mediated knockdown for epistatic analysis of flies carrying *opa1*^i^, *catalase*^i^, *sod2*^i,^ and *erk*^i^ in the FRT40A and *drp1*^KG^ FRT40A shown above were crossed to *hs*-FLP; Gal80 FRT40A/CyO; *tub*-Gal4,UAS-CD8-GFP/TM6, and appropriate flies were selected to perform heat shocks and assess the phenotypes in clones after 10 days.

### Immunostaining of the *Drosophila* ovaries

*Drosophila* ovaries were dissected 10 days post-heat shock in a serum-free Schneider’s insect medium and ovarioles were separated using fine needles. The dissected ovaries were washed in fresh Schneider’s insect medium to remove debris released during dissection and fixed using freshly prepared 4% PFA in 1XPBS for 20-25 min followed by three washes of in 1X PBS containing 0.3% Triton X-100 (T8787-Sigma) (further called 0.3% PBST) for 10 min each. The ovaries were transferred to a blocking solution containing 2% BSA (MB083, HIMEDIA) dissolved in 0.3% PBST for 1 hr and incubated with primary antibodies for 16-18 hours followed by three 10 min washes with 0.3% PBST. The secondary antibodies, dissolved in 0.3% PBST, were added for 2 hr. The ovaries were washed three times in 0.3% PBST for 5 min each. The second wash of 0.3% PBST contained a Hoechst 33258 (H3569, ThermoFisher), with 1:1000 dilution. The ovarioles were separated and mounted on a glass slide with Slowfade (S36937, ThermoFisher). The primary antibodies used are 1:500 chicken GFP (A10262, ThermoFisher), 1:500 rabbit aPKC, 1:500 mouse aPKC, 1:10 mouse Dlg (4F3, DSHB), 1:10 mouse Hindsight (1G9, DSHB), 1:200 rabbit dpERK (4370, CST), 1:200 mouse ATPβ (ab14730, Abcam). The secondary antibodies used at a 1:1000 dilution are anti-chicken 488 (A11039, ThermoFisher), anti-mouse 568 (A11004, ThermoFisher), anti-mouse 633 (A21050, ThermoFisher), anti-mouse 647 (A21235, ThermoFisher), anti-rabbit 568 (A11011, ThermoFisher), anti-rabbit 633 (A21070, ThermoFisher), anti-rabbit 647 (A21245, ThermoFisher).

### MitoSOX staining for mitochondrial ROS

The MitoSOX (M36008, ThermoFisher) dye gets oxidized by mitochondrial superoxides and gives fluorescence (Robinson et al., 2006). The ovaries were dissected and washed in Schneider’s medium. The ovaries were immersed in Schneider’s medium containing MitoSOX dye with a final concentration of 5 min wash with Schneider’s medium (Parker et al., 2017). The ovaries were immediately transferred to a glass slide and mounted using Schneider’s medium for the imaging.

### Data quantification and Image analysis

#### Identification of stages of development in ovarioles

The stage identification was performed by measuring the surface area of the middle stack of the egg chambers (Jia et al., 2016). Images were quantified with the help of Fiji and ImageJ (Schindelin et al., 2012; Schneider et al., 2012).

### Quantification of multilayering

The estimation of the number of layers of FCs was performed for PFCs in controls and mutants. The rows of cells per clone were counted in the mCD8-GFP positive clones to report the phenotype of multilayering. The analysis of multilayering of *drp1*^KG^ used for comparison in other genotypes remained the same.

### Quantification of mitochondrial morphology

To evaluate the mitochondrial morphology, the Streptavidin or ATP-β staining of the mitochondria in the GFP positive follicular cell clones and GFP negative neighboring cells were compared visually. Streptavidin or ATP-β staining in the follicle cells that are more punctate was used to characterize the fragmented mitochondria. The Compactness of the mitochondrial staining and a punctate appearance from the ATP-β staining was used to distinguish between mitochondria that were more and less clustered. Furthermore, the follicle cells’ mitochondrial morphology, which lacked any discernible variations, was classified as intermediate. From the observations, the percentage of egg chambers with the respective mitochondrial morphology was estimated.

### Estimation of numbers of cells containing aPKC and Hnt

For the analysis of cells containing polarity protein aPKC on the apical membrane, we estimated the numbers of FCs per clone deficient of aPKC at the apical membrane in the mCD8-GFP positive clones and expressed them as a percentage of total cells in the clone. To quantitate the appearance of the aPKC on the apical membrane in epistasis experiments, the percentage of PFCs containing apical aPKC was counted in the layer of cells adjacent to the oocyte. For analysis of Notch signaling mediated differentiation, we counted a percentage of the egg chambers based on the presence of the Hnt for the comparisons among genotypes.

### Quantification of dpERK

For the analysis of relative dpERK fluorescence in different genotypes, we marked a region of interest (ROI) around the clone expressing mCD8-GFP and neighboring FCs. A background ROI was used to subtract the noise from the actual signal. Appropriate thresholding was applied to quantitate the dpERK in cells. The ratio of the thresholded intensity value of dpERK from the clone to their neighboring cell was computed.

### n values and statistics

Each data point (n) represents quantification from a distinct clone in different egg chambers. A minimum number of 8 egg chambers are used for the statistics. The data for each genotype is shown with its respective mean and SEM. The statistical test applied for two groups is Student’s t-test and for more than two groups is One-way analysis of variance with Tukey’s multiple comparison test.

## Results

### Drp1 depleted PFCs are present in multiple layers in the endocycling stage, and this defect is partially suppressed by the depletion of Opa1

To analyze the organization of *Drosophila* epithelial FCs depleted of mitochondrial fission protein Drp1, we generated MARCM clones (see materials and methods) with the transposon tagged null mutant *drp1*^KG03815^ (*drp1*^KG^) (Mitra et al., 2012; Rikhy et al., 2007; Tomer et al., 2018). Drp1 depleted FCs are marked by the expression of mCD8-GFP. Ovaries containing FC clones depleted of Drp1 were stained for nuclear DNA and mitochondria. As expected, Drp1 depletion led to mitochondrial clustering in FCs (Fig. S1A,B) similar to the previous analysis in neuroblasts, neurons, FCs, and spermatocytes (Aldridge et al., 2007; Dubal et al., 2022; Mitra et al., 2012; Tomer et al., 2018). Drp1 depleted FC clones were present in a monolayer at the mitotic stage egg chambers (Fig. 1B). PFC clones at the endocycling stage depleted of Drp1 were present in multiple layers (Fig. 1C). The number of cell layers per clone in the *drp1*^KG^ depleted PFCs varied from 1 to 6 with the highest frequency of 3 layers (Fig. S1C).

Mitochondrial clustering in Drp1-depleted cells requires the activity of mitochondrial fusion proteins Opa1 and Marf (Dubal et al., 2022; Sessions et al., 2022). We depleted Opa1 using an shRNA to inhibit mitochondrial clustering in Drp1-depleted FCs. Mitochondria were fragmented in Opa1 depleted FCs (Fig. S1A,B). Even though the mitochondria were clustered in the *drp1*^KG^;*opa1*^i^ combination, the cluster appeared to be decreased in compaction as compared to *drp1*^KG^ alone (Fig. S1A,B). FCs depleted of Opa1 were present in a single layer similar to controls in the mitotic and endocycling stages (Fig. 1B, C). The *drp1*^KG^;*opa1*^i^ combination showed the presence of PFCs in multiple layers, however, the extent of multilayering was reduced as compared to *drp1*^KG^ (Fig. 1C, D). In addition to estimating the number of layers in each clone, we quantified the clone area in different genotypes as a readout of the extent of multilayering. Consistent with the decrease in multilayering, we found a significant reduction in the clone area of *drp1*^KG^;*opa1*^i^ PFCs as compared to *drp1*^KG^ alone (Fig. 1D). In summary, Drp1 depleted PFCs were present in multiple layers at the endocycling stage and additional depletion of Opa1 partially alleviated the extent of multilayering.

### Drp1 depleted FCs show loss of apical polarity protein aPKC

Depletion of polarity proteins aPKC, Baz, Crumbs, Scribble, Dlg, and Lgl leads to the distribution of FCs in multiple layers (Baum and Perrimon, 2001; Benton and St Johnston, 2003; Bergstralh et al., 2013; Bilder et al., 2000; Dent et al., 2019; Fletcher et al., 2012; Khoury and Bilder, 2020; Luo et al., 2016; Moreira et al., 2019; Romani et al., 2009; Sun and Deng, 2005; Ventura et al., 2020; Wang et al., 2021; Woolworth et al., 2009). The apical protein aPKC phosphorylates several other polarity proteins including Dlg, Bazooka, Crumbs, Lgl, and coordinates epithelial cell polarization (Betschinger et al., 2003; Golub et al., 2017; Morais-de-Sá et al., 2010; Plant et al., 2003; Sotillos et al., 2004). In addition, aPKC also maintains a single layer of FCs by inhibiting actomyosin contractility (Osswald et al., 2022). We assessed the distribution of apical aPKC and lateral Dlg in *drp1*^KG^ PFC clones in endocycling stages and FC clones in mitotic stages. The aPKC levels were reduced from PFCs depleted of Drp1 located adjacent to the oocyte in the endocycling stage (Fig. 2A). At earlier mitotic stages, despite being arranged in a monolayer similar to controls, the *drp1*^KG^ FCs also showed depletion of aPKC (Fig. 2C). This aPKC depletion was suppressed in *drp1*^KG^;*opa1*^i^ combination in the endocycling PFCs adjacent to the oocyte (Fig. 2A, C) and the mitotic stage FCs (Fig. 2B, D). We further assessed if Dlg would spread into the apical domain in *drp1*^KG^ FCs due to the depletion of aPKC (Schmidt and Peifer, 2020). Dlg remained in the lateral domain in PFCs adjacent to oocyte even at the endocycling stage and in the mitotic stage FCs depleted of Drp1 (Fig. S2A, B). However, the FCs in the mitotic stage appeared constricted in the majority of the Drp1 depleted FCs (Fig. S2A, B). Since aPKC is known to inhibit the contractility of the actomyosin cytoskeleton (Osswald et al., 2022), its depletion likely contributes to the change in the shape of FCs in mitotic stages and further occurrence of PFCs in multiple layers on Drp1 depletion.

**Figure 2:**
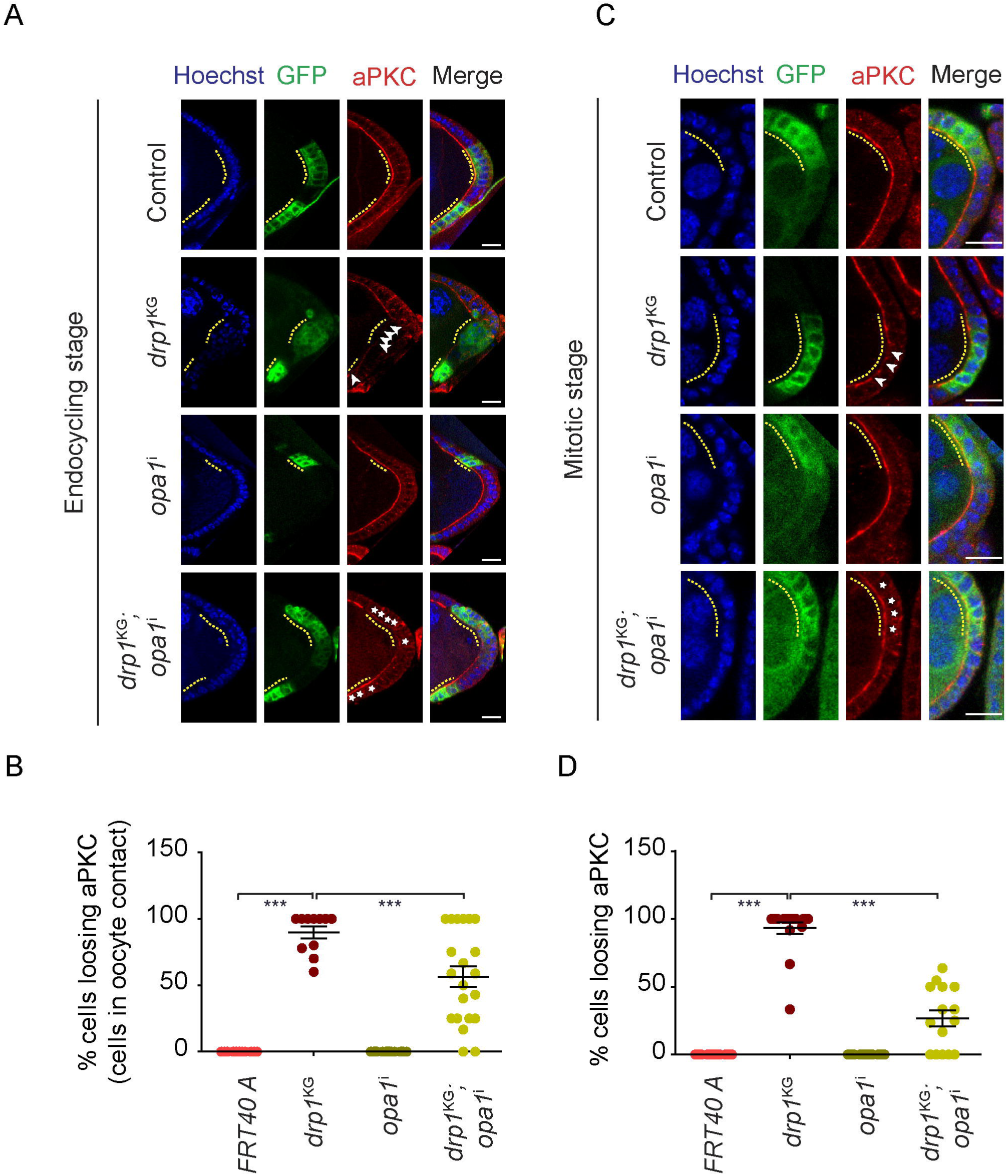
Drp1 depleted FC clones show aPKC loss in endocycling and mitotic stages and this defect is decreased in *drp1*^KG^;*opa1*^i^. **(A-D)** Representative images showing aPKC (red) in PFC clones from *drp1*^KG^ and *drp1*^KG^;*opa1*^i^ at the endocycling stage **(A)**. The graph shows the percentage of cells per PFC clone adjacent to the oocyte losing aPKC in *drp1*^KG^ and *drp1*^KG^;*opa1*^i^ at the endocycling stage **(B).** Representative images showing aPKC (red) in *drp1*^KG^ and *drp1*^KG^;*opa1*^i^ in mitotic stage FCs **(C)**. The graph shows the percentage of cells per FC clone in mitotic stages losing aPKC in *drp1*^KG^ and *drp1*^KG^;*opa1*^i^ **(D)**. mCD8-GFP (green) expressing FC clones of the indicated genotype are marked with dashed yellow outlines. The nucleus (blue) is stained with Hoechst. Arrowheads show defective in FCs. Asterisks mark FCs that show the presence of aPKC in double mutants. The graphs show data in Mean±SEM. Each data point (n) in the endocycling and mitotic stages represents the percentage of cells losing aPKC in a single clone of a separate egg chamber. The n values for the genotypes control FRT40A, *drp1*^KG^, *opa1*^i^, and *drp1*^KG^;*opa1*^i^ is given as (n, N; n is the number of clones, N is the number of independent replicates) being (n=11,11,22,21 and N=3,5,3,4 respectively) for endocycling stages and (n=17,17,21,15 and N=3,5,3,4 respectively) for mitotic stages. The statistical test performed is one-way ANOVA with Tukey’s multiple comparisons. ns=non-significant, * P<0.05, ** P<0.01, *** P<0.001, Scale bar=10μm.

### ROS increase on depletion of Opa1 and ROS scavenging proteins leads to mitochondrial fragmentation

Opa1 depletion partially alleviated the aPKC loss of Drp1 deficient FCs (Fig. 2). Loss of Opa1 function has been previously shown to increase mitochondrial ROS and overexpression of Opa1 along with the ATP synthase complex oligomerization reduced mitochondrial ROS (Jang and Javadov, 2020; Quintana-Cabrera et al., 2021; Tang et al., 2009). Mitochondrial ROS (mtROS) was found to be elevated in the Opa1-depleted eye cells in *Drosophila (Yarosh et al., 2008)*. On the other hand, ROS was also found to be decreased on Drp1 depletion in *Drosophila* embryos (Chowdhary et al., 2020). We stained live ovaries containing control or mutant clones with a fluorescent dye mitoSOX for the detection of mitochondrial superoxides. Depletion of Opa1 led to an increase in the mitoSOX fluorescence as compared to their neighboring control FCs (Fig. 3A). ROS levels are regulated by antioxidant enzymes such as superoxide dismutase 1 (SOD1), SOD2, catalase, and glutathione peroxidase. We, therefore, tested if an increase in ROS was seen when we depleted mitochondrial SOD2 (Celotto et al., 2012; Kirby et al., 2002; Mukherjee et al., 2011) and catalase using shRNA expression. The fluorescence intensity of mitoSOX was higher in *sod2*^i^ and *catalase*^i^ FC clones as compared to the neighboring cells (Fig. 3A). Additionally, SOD2 depletion has also been shown to affect mitochondrial dynamics leading to fragmentation in delaminating cells during *Drosophila* dorsal closure (Muliyil and Narasimha, 2014). We, therefore, assessed mitochondrial morphology in *sod2*^i^ and *catalase*^i^ FC clones by immunostaining with an antibody against Complex V. Mitochondria were found to be punctate in appearance in FCs depleted of *sod2*^i^ and *catalase*^i^ as compared to neighboring control cells (Fig. 3B). The mitochondrial morphology in *drp1*^KG^;*sod2*^i^ and in *drp1*^KG^;*catalase*^i^ showed mitochondrial clustering but the compaction was decreased compared to *drp1*^KG^ (Fig. S3A, B). Mitochondria were typically distributed to one side in multilayered PFCs depleted of Drp1 (Fig. S3B). Mitochondria were present all around in the nucleus consistent with fragmentation in the *drp1*^KG^;*sod2*^i^ and in *drp1*^KG^;*catalase*^i^ combinations as compared to *drp1*^KG^ (Fig. S3B). In summary, we found that the depletion of Opa1 led to increased ROS and mitochondrial fragmentation similar to SOD2 and catalase depletion.

**Figure 3:**
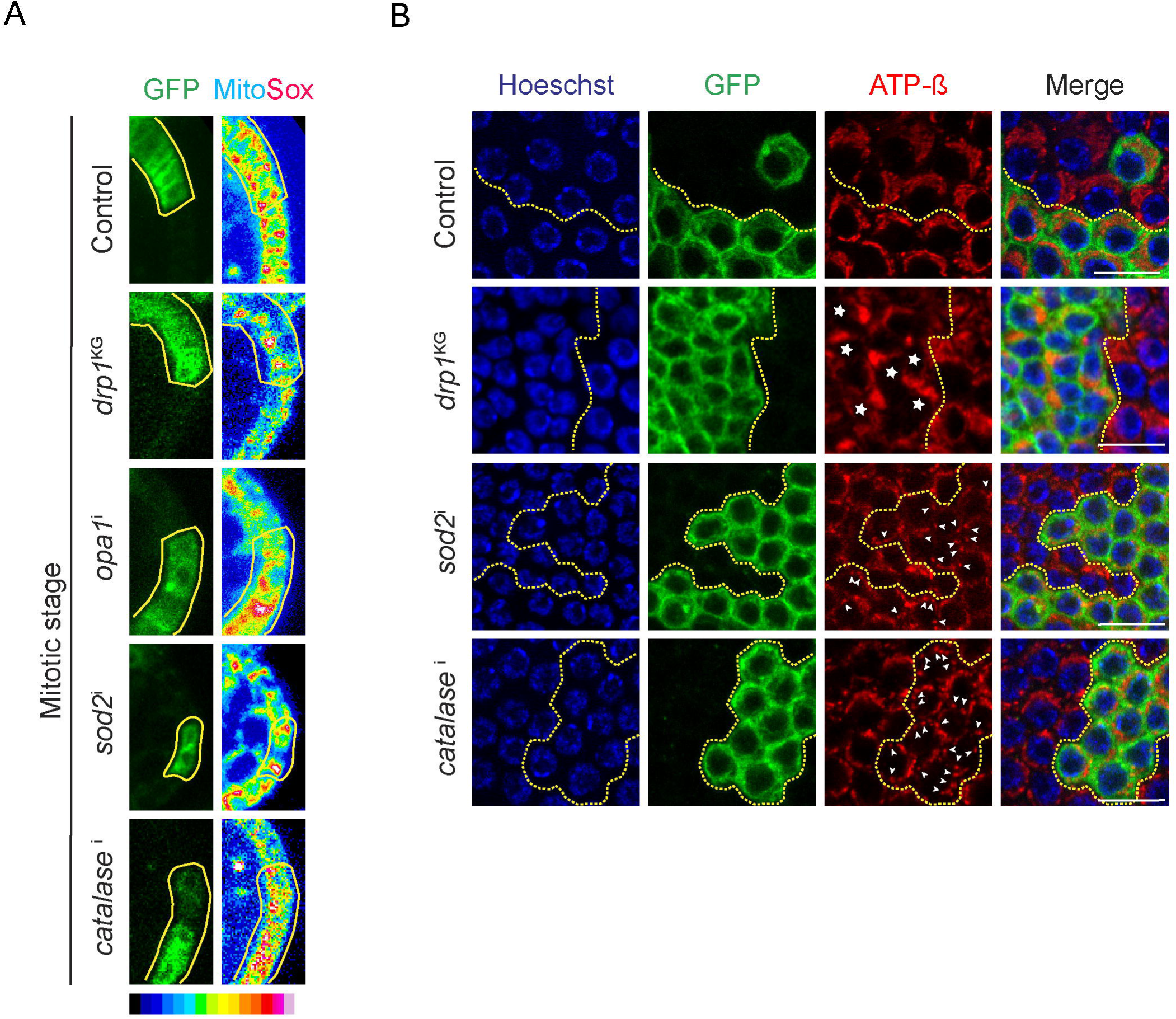
Mitochondrial ROS and fragmentation is increased on depletion of ROS scavengers. **(A-B)** Representative images showing the fluorescence intensity in a rainbow scale (red pixels are higher intensity and blue pixels are lower intensity) of MitoSOX in control FRT40A, *drp1*^KG^, *opa1*^i^, *sod2*^i^, and *catalase*^i^ **(A)** M. Mitochondria stained with ATP (red) are shown in representative images from control FRT40A (100% intermediate, n=21), *drp1*^KG^ (92% clustered, n=26), *sod2*^i^ (82% fragmented, n=28), and *catalase*^i^ (79% fragmented, n=14) **(B)**. mCD8-GFP (green) expressing FC clones of the indicated genotype are marked with solid or dashed yellow outlines. Asterisks mark clustered mitochondria in surface view. Arrowheads mark the punctate mitochondria in surface view. The nucleus (blue) is stained with Hoechst. Scale bar=10μm.

### ROS increase in Drp1 depleted PFCs shows an alleviation of polarity defects and loss of aPKC

ROS alters the activity and distribution of protein kinases by oxidation (Corcoran and Cotter, 2013). Protein kinase C (PKC) is activated by mtROS in human peritoneal mesothelial cells (HPMC), and PKC can in turn further enhance the ROS, to continue the activation cycle (Lee et al., 2004). ROS plays a significant role in the regulation of PKC activation. Since *drp1*^KG^;*opa1*^i^ FCs showed alleviation of multilayering and loss of aPKC defect as compared to *drp1*^KG^ alone, we tested if depletion of ROS scavengers in Drp1 depleted FCs improved their organization and aPKC levels. The percentage of the egg chambers with 3 or more layers of FCs in *drp1*^KG^;*sod2*^i^ and *drp1*^KG^;*catalase*^i^ mutant clones was reduced as compared to *drp1*^KG^ (Fig. S4A, C). There was a significant reduction in the area of the clone in *drp1*^KG^;*sod2*^i^ and *drp1*^KG^;*catalase*^i^ as compared to *drp1*^KG^ (Fig. S4B, D). The levels of aPKC in endocycling PFCs adjacent to the oocyte (Fig 4A, B) and mitotic stage FC clones (Fig. 4C, D) of *drp1*^KG^;*sod2*^i^ and *drp1*^KG^;*catalase*^i^ increased on the apical membrane as compared to *drp1*^KG^ alone. This increase in aPKC was similar to that found in the double mutant *drp1*^KG^;*opa1*^i^ clones (Fig. 2A-D). Thus, a partial recovery of multilayering and aPKC levels observed in *drp1*^KG^;*sod2*^i,^ and *drp1*^KG^;*catalase*^i^ suggested an important role of ROS in regulating mitochondrial morphology and distribution of apical polarity protein aPKC in follicle epithelial cells.

**Figure 4:**
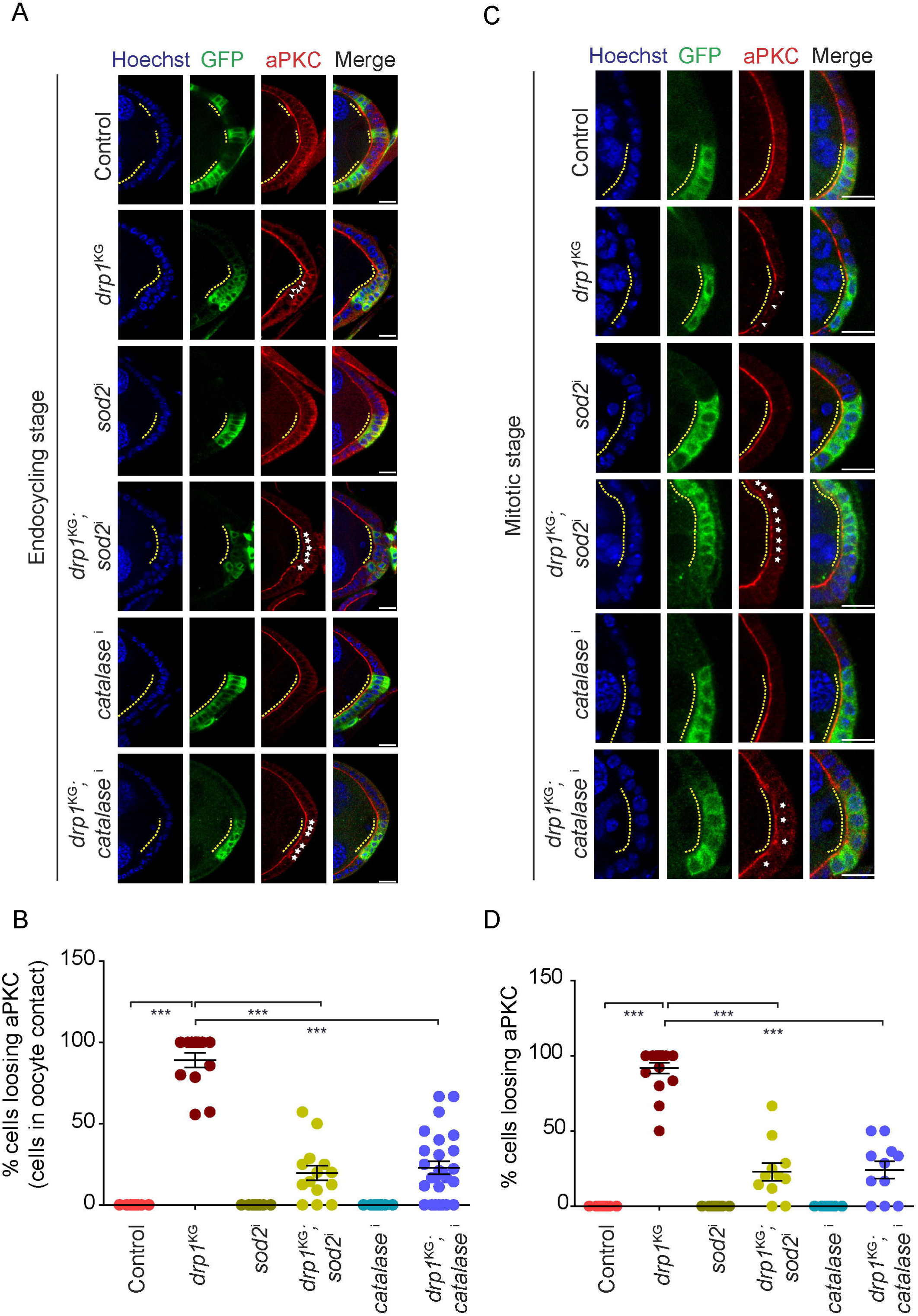
Drp1 depleted FC clones show loss of aPKC in endocycling and mitotic stages and this is partially recovered on additional depletion of ROS scavengers. **(A-D)** Representative images showing aPKC (red) staining in control FRT40A, *drp1*^KG^, *sod2*^i^, *drp1*^KG^;*sod2*^i^, *catalase*^i^ and *drp1*^KG^;*catalase*^i^ PFC clones (A). The graph shows quantification of the percentage of cells losing aPKC in each PFC clone adjacent to the oocyte in control FRT40A, *drp1*^KG^, *sod2*^i^, *drp1*^KG^;*sod2*^i^, *catalase*^i^ and *drp1*^KG^;*catalase*^i^ (B). Representative images showing aPKC (red) staining in control FRT40A, *drp1*^KG^, *sod2*^i^, *drp1*^KG^;*sod2*^i^, *catalase*^i^ and *drp1*^KG^;*catalase*^i^ FC clones in mitotic stages (C). The graph shows the quantification of the percentage of cells losing aPKC in each FC clone in mitotic stages in control FRT40A, *drp1*^KG^, *sod2*^i^, *drp1*^KG^;*sod2*^i^, *catalase*^i^ and *drp1*^KG^;*catalase*^i^ (D) mCD8-GFP (green) expressing FC clones of the indicated genotype are marked with dashed yellow outlines. The nucleus (blue) is stained with Hoechst. Arrowheads mark FCs deficient in aPKC. Asterisks mark FCs that show the presence of aPKC in double mutants. The graphs show data in Mean±SEM. Each data point (n) in the endocycling and mitotic stages represents the percentage of cells losing aPKC in a single clone of a separate egg chamber. The n values of the genotypes control FRT40A, *drp1*^KG^, *sod2*^i^, *drp1*^KG^;*sod2*^i^, *catalase*^i,^ and *drp1*^KG^;*catalase*^i^ given as (n, N; n is the number of clones, N is the number of independent replicates) (n=12,13,9,14,13,26 and N=3,5,3,3,3,3 respectively) for endocycling stages and (n=17,17,26,11,15,11 and N=3,4,3,3,3,3 respectively) for mitotic stages. The statistical test performed is One way ANOVA with Tukey’s multiple comparisons. ns=non-significant, * P<0.05, ** P<0.01, *** P<0.001, Scale bar=10μm.

### ERK accumulation in Drp1 depleted FCs is suppressed by additional depletion of Opa1 and ROS scavengers

EGFR signaling leads to the activation of ERK by phosphorylation (González-Reyes and St Johnston, 1998). Previous studies have shown that EGFR signaling-driven accumulation of doubly phosphorylated ERK (dpERK) increases in the cytoplasm in Drp1 depleted PFCs (Tomer et al., 2018). A decrease in EGFR signaling in pre-FCs is important for the recruitment of aPKC and the formation of the apical domain (Castanieto et al., 2014). We have shown that aPKC is decreased in Drp1 depleted cells (Fig. 2,4). Since ROS increase in Drp1 depleted FCs led to an increase in aPKC presence at the apical membrane, we assessed if this was coincident with a loss of accumulation of dpERK. For this, we quantified the change in dpERK on inhibiting Opa1 and ROS scavengers in Drp1 depleted FCs. We found that there was a reduction in the accumulation of dpERK in mitotic stage FCs (Fig. 5A, B) and PFCs adjacent to the oocyte in endocycling stages (Fig. 5C, D) in the *drp1*^KG^;*opa1*^i^, *drp1*^KG^;*sod2*^i^ and *drp1*^KG^;*catalase*^i^ combination as compared to *drp1*^KG^ alone. The increase in aPKC in Drp1 depleted FCs on additionally depleting Opa1 and ROS scavengers is accompanied by a loss of dpERK in the cytoplasm.

**Figure 5:**
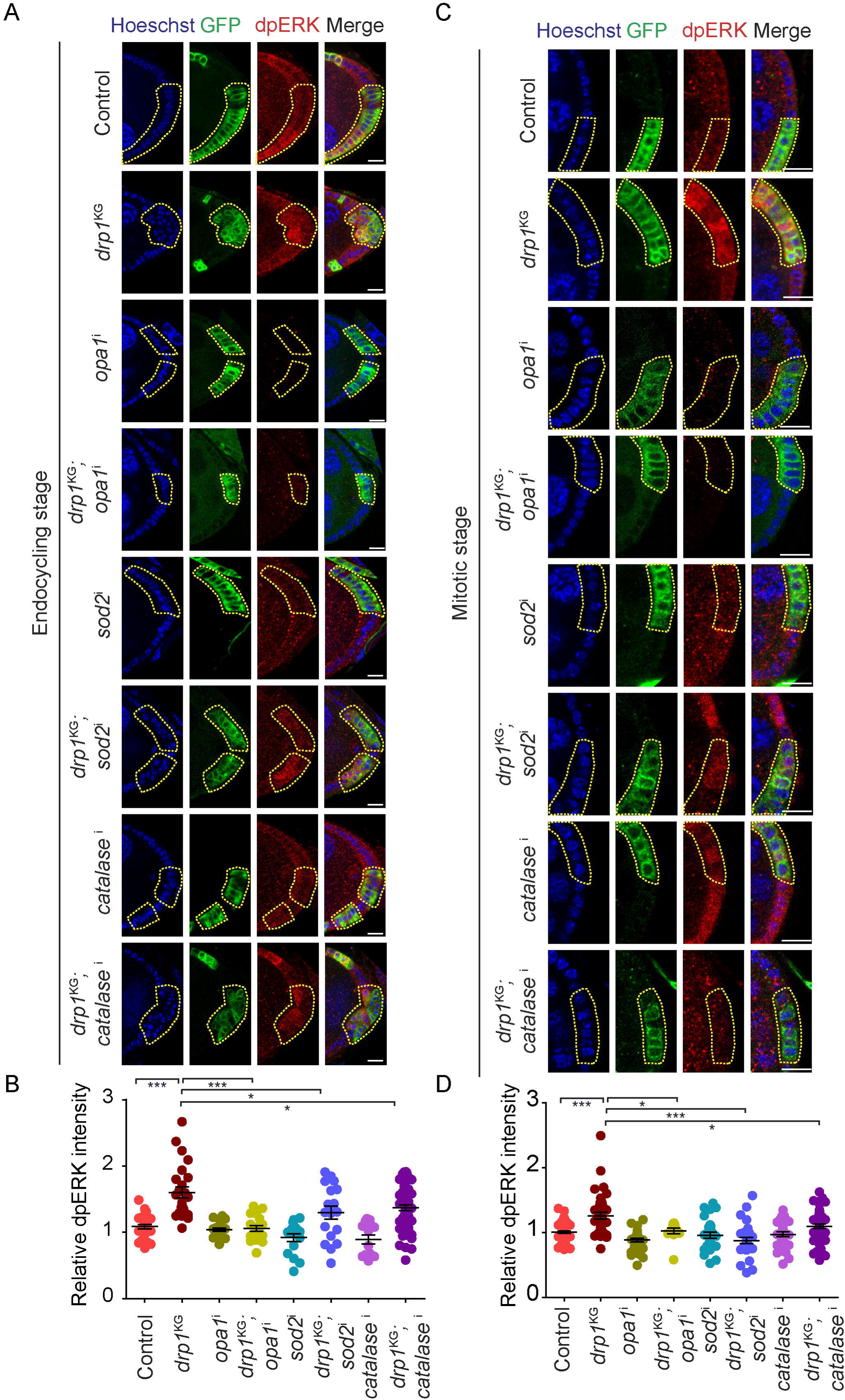
Increase in dpERK in Drp1 depleted clones is rescued on additional depletion of Opa1 and ROS scavengers. **(A-D)** Representative images showing dpERK (red) in control FRT40A, *drp1*^KG^, *opa1*^i^, *drp1*^KG^;*opa1*^i^, *sod2*^i^, *drp1*^KG^;*sod2*^i^, *catalase*^i,^ and *drp1*^KG^;*catalase*^i^ PFC clones at endocycling stages **(A)**. Representative images showing dpERK (red) in control FRT40A, *drp1*^KG^, *opa1*^i^, *drp1*^KG^;*opa1*^i^, *sod2*^i^, *drp1*^KG^;*sod2*^i^, *catalase*^i,^ and *drp1*^KG^;*catalase*^i^ FC clones at mitotic stages **(B)**. The graph shows the quantification of relative dpERK fluorescence compared to neighboring control cells in control FRT40A, *drp1*^KG^, *opa1*^i^, *drp1*^KG^;*opa1*^i^, *sod2*^i^, *drp1*^KG^;*sod2*^i^, *catalase*^i,^ and *drp1*^KG^;*catalase*^i^ PFC clones at endocycling stage **(C)**. The graph shows the quantification of relative dpERK fluorescence compared to neighboring control cells in control FRT40A, *drp1*^KG^, *opa1*^i^, *drp1*^KG^;*opa1*^i^, *sod2*^i^, *drp1*^KG^;*sod2*^i^, *catalase*^i,^ and *drp1*^KG^;*catalase*^i^ FC clone at mitotic stages **(D).** mCD8-GFP (green) expressing FC clones of the indicated genotype are marked with dashed yellow outlines. The nucleus (blue) is stained with Hoechst. The data of each genotype is presented with its respective mean and SEM. The graphs show data in Mean±SEM. Each data point (n) in the endocycling and mitotic stages represents the relative fluorescence intensity of dpERK from a single clone from a separate egg chamber. The n values for genotypes control FRT40A, *drp1*^KG^, *opa1*^i^, *drp1*^KG^;*opa1*^i^, *sod2*^i^, *drp1*^KG^;*sod2*^i^, *catalase*^i^, and *drp1*^KG^;*catalase*^i^ are given as (n, N; n is the number of clones, N is the number of independent replicates) (n=25,24,20,18,16,18,14,55 and N=4,4,3,4,3,3,3,6 respectively) for endocycling stages and (n=52,40,31,12,27,30,30,47 and N=5,4,2,4,3,3,2,5 respectively) for mitotic stages. ns=non-significant, * P<0.05, ** P<0.01, *** P<0.001, Scale bar=10μm.

### ERK loss in Drp1 depleted FCs leads to alleviation of the polarity and aPKC defects

Since ERK decreased in Drp1 depleted FCs on the additional expression of RNAi against Opa1 and ROS scavengers, we tested if ERK depletion would lead to a change in polarity and apical aPKC accumulation. As expected, ERK RNAi (*erk*^i^) expression led to decreased dpERK levels in the FCs of both endocycling and mitotic stages in *drp1*^KG^ (Fig. S5A-D). The percentage of the egg chambers showing 3 or more layers in *drp1*^KG^;*erk*^i^ FCs was reduced as compared to *drp1*^KG^ alone (Fig. S6A). There was a significant reduction in the area of the clone in *drp1*^KG^;*erk*^i^ as compared to *drp1*^KG^ (Fig. S6B). We observed that, unlike *drp1*^KG^, aPKC was present in *drp1*^KG^;*erk*^i^ (Fig. 6A-D) in mitotic FCs and endocycling PFCs adjacent to the oocyte. A decrease in ERK downstream of EGFR signaling led to the alleviation of polarity defects in Drp1 depleted cells.

**Figure 6:**
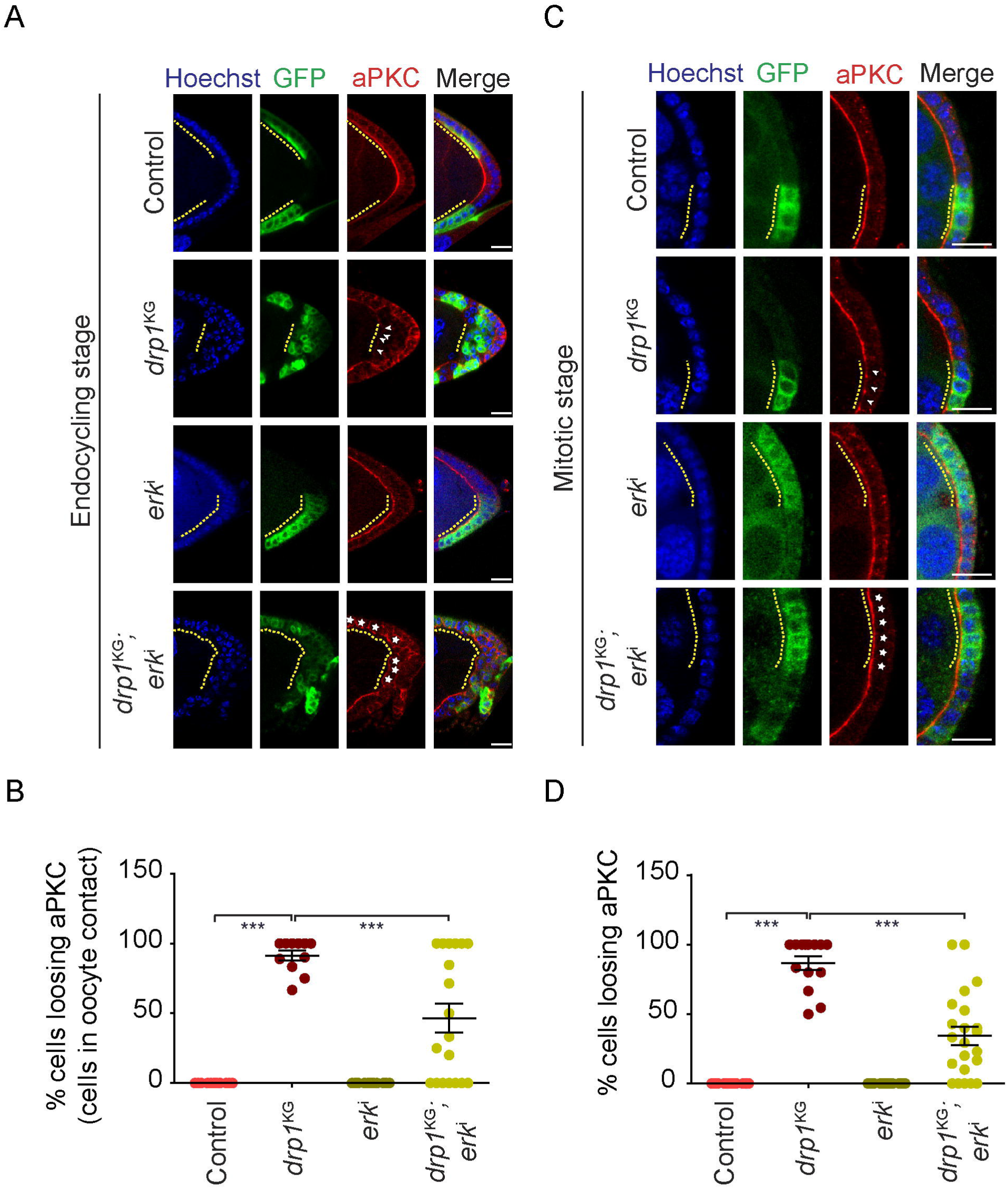
aPKC depletion in Drp1 depleted FC clones recovers on additional depletion of ERK. **(A-D)** Representative images showing aPKC (red) in control FRT40A, *drp1*^KG^, *erk*^i^, and *drp1*^KG^;*erk*^i^ PFC clones at endocycling stages **(A)**. Representative images showing aPKC (red) in control FRT40A, *drp1*^KG^, *erk*^i^, and *drp1*^KG^;*erk*^i^ FC clones at mitotic stages **(B).** The graph shows the quantification of the percentage of cells losing aPKC in each PFC clone adjacent to the oocyte in control FRT40A, *drp1*^KG^, *erk*^i^, and *drp1*^KG^;*erk*^i^ at endocycling stages **(C)**. The graph shows the quantification of the percentage of cells losing aPKC in each FC clone in control FRT40A, *drp1*^KG^, *erk*^i^, and *drp1*^KG^;*erk*^i^ at mitotic stages **(D).** mCD8-GFP (green) expressing FC clones of the indicated genotype are marked with dashed yellow outlines. The nucleus (blue) is stained with Hoechst. Arrowheads mark FCs deficient in aPKC. Asterisks mark FCs that show the presence of aPKC in double mutants. The graphs show data in Mean±SEM. Each data point (n) in the endocycling and mitotic stages represents the percentage of cells losing aPKC in a single clone of a separate egg chamber. The n values for the genotypes control FRT40A, *drp1*^KG^, *erk*^i,^ and *drp1*^KG^;*erk*^i^ is given as (n,N; n is the number of clones, N is the number of independent replicates) (n=9,11,10,19 and N=3,5,3,4 respectively) for endocycling stages and (n=23,14,12,22 and N=5,5,3,3 respectively) for mitotic stages. ns=non-significant, * P<0.05, ** P<0.01, *** P<0.001. Scale bar=10μm.

Even though Drp1 depleted FCs showed an increase in dpERK, this increase occurred in the cytoplasm and there was a loss of oocyte patterning depicted by oocyte migration to the dorso-anterior location (Mitra et al., 2012; Tomer et al., 2018). We found oocyte remained in the posterior location in *drp1*^KG^;*opa1*^i^, *drp1*^KG^;*sod2*^i^, and *drp1*^KG^;*catalase*^i^ at a frequency similar to *drp1*^KG^ (Fig. S7). These data showed that even though the cytoplasmic accumulation of dpERK was reduced on depletion of Opa1 and ROS scavengers in Drp1 depleted FCs, the defect in oocyte migration and patterning still remained.

### The Notch signaling defect in Drp1 deficient FCs is suppressed by the depletion of ROS scavengers

Notch signaling activation promotes the transition of the mitotic stage to the endocycling stage in FCs during oogenesis by the expression of transcription factor Hindsight (Hnt) (Kim-Yip and Nystul, 2018; López-Schier and St Johnston, 2001; Ruohola et al., 1991). Drp1 depletion leads to the loss of Hnt due to altered mitochondrial activity and accumulation of dpERK (Mitra et al., 2012; Tomer et al., 2018). We further assessed if the increase in ROS on depletion of ROS scavenging enzymes in Drp1 deficient FCs could lead to a suppression of the defect in Notch signaling activation. We found that the depletion of Opa1, SOD2, Catalase, and ERK showed the activation of Hnt similar to controls. The *drp1*^KG^;*opa1*^i^, *drp1*^KG^;*sod2*^i^, *drp1*^KG^;*catalase*^i^, and *drp1*^KG^;*erk1*^i^ combinations, showed expression of Hnt unlike *drp1*^KG^ (Fig. 7). These data showed that the loss of Notch signaling mediated differentiation was alleviated by the increase in ROS in FCs depleted for Drp1.

**Figure 7:**
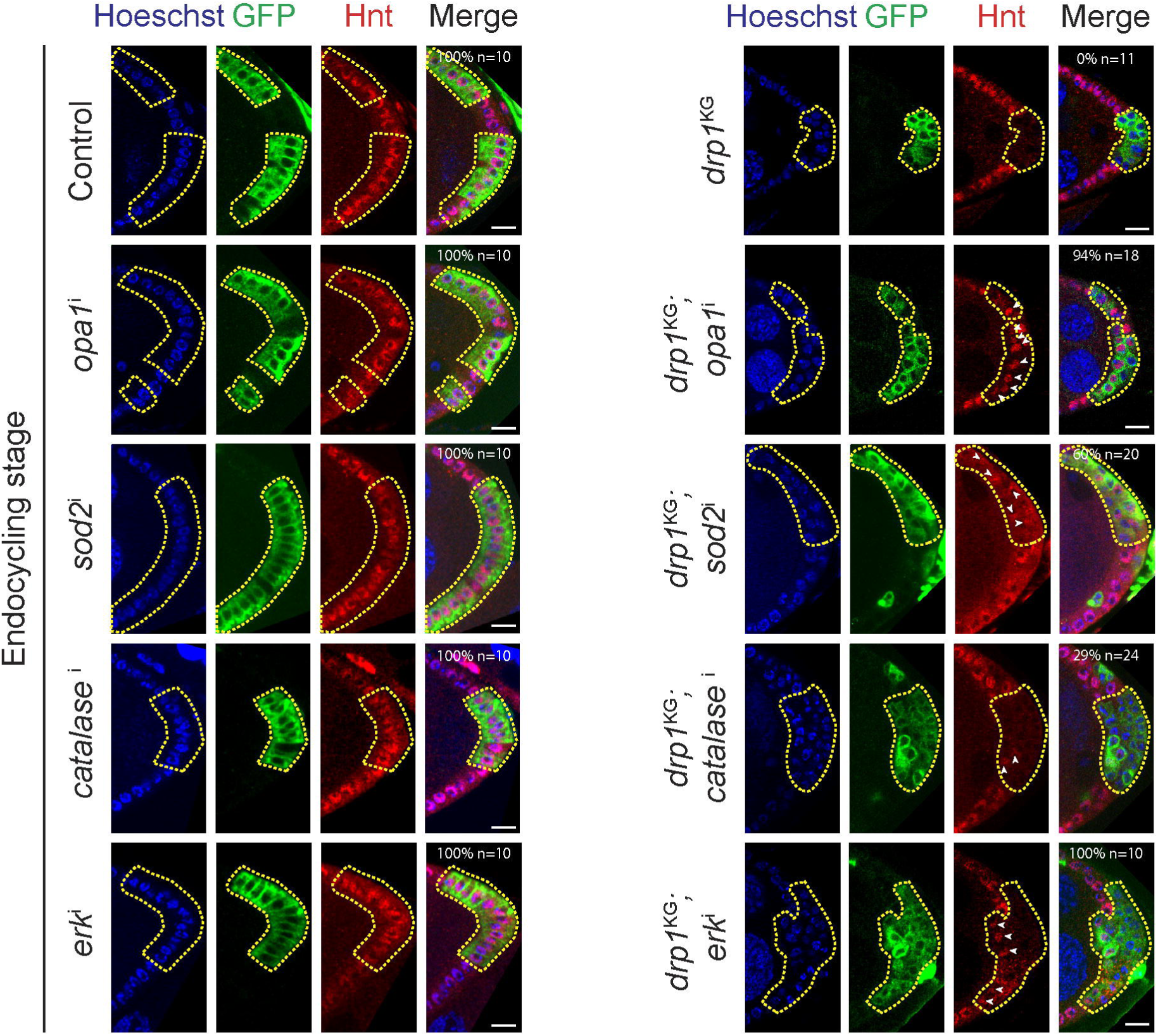
Notch differentiation defects in Drp1 depleted PFC clones are suppressed on additional depletion of Opa1, ROS scavengers, and ERK. Representative images showing Hnt (red) in PFCs in the genotypes control FRT40A (100% show Hnt, n=10), *opa1*^i^ (100%,n=10), *sod2*^i^ (100%,n=10), *catalase*^i^ (100%,n=10) and *erk*^i^ (100%,n=10) on the left side and *drp1*^KG^ PFCs is (0%,n=11), *drp1*^KG^;*opa1*^i^ (94%,n=18), *drp1*^KG^;*sod2*^i^ (60%,n=20), *drp1*^KG^;*catalase*^i^ (29%,n=24), *drp1*^KG^;*erk1*^i^ (100%,n=10) on the right side. % denotes the percentage of egg chambers with PFC clones that express Hnt in PFC clones. mCD8-GFP (green) expressing FC clones of the indicated genotype are marked with dashed yellow outlines. The nucleus (blue) is stained with Hoechst. Arrowheads mark PFCs showing the presence of Hnt. The numbers in each image represent the percentage of PFC clones showing the presence of Hnt. Scale bar=10μm.

In summary, our results show that an increase in ROS can alleviate the mitochondrial fusion, aPKC, and Notch signaling defect in Drp1 depleted FCs.

## Discussion

In this study, we show that mitochondrial fission by Drp1 regulates apical polarity and signaling via mitochondrial morphology and ROS in *Drosophila* follicle epithelial cell differentiation. We find that appropriate ROS levels regulated by mitochondrial fragmentation are important for the interaction between activated ERK and aPKC in mitotic stage FCs (Fig. 8). ROS also plays a key role in the interaction between EGFR and Notch signaling to differentiate PFCs during oogenesis (Fig. 8). We discuss our results in the following three contexts: 1] the role of depletion of aPKC in causing multilayering of PFCs, 2] the function of ROS in regulating the onset of apical polarity and 3] the interaction of mitochondrial dynamics, ROS, and activation of signaling pathways in epithelial polarity formation and differentiation.

**Figure 8:**
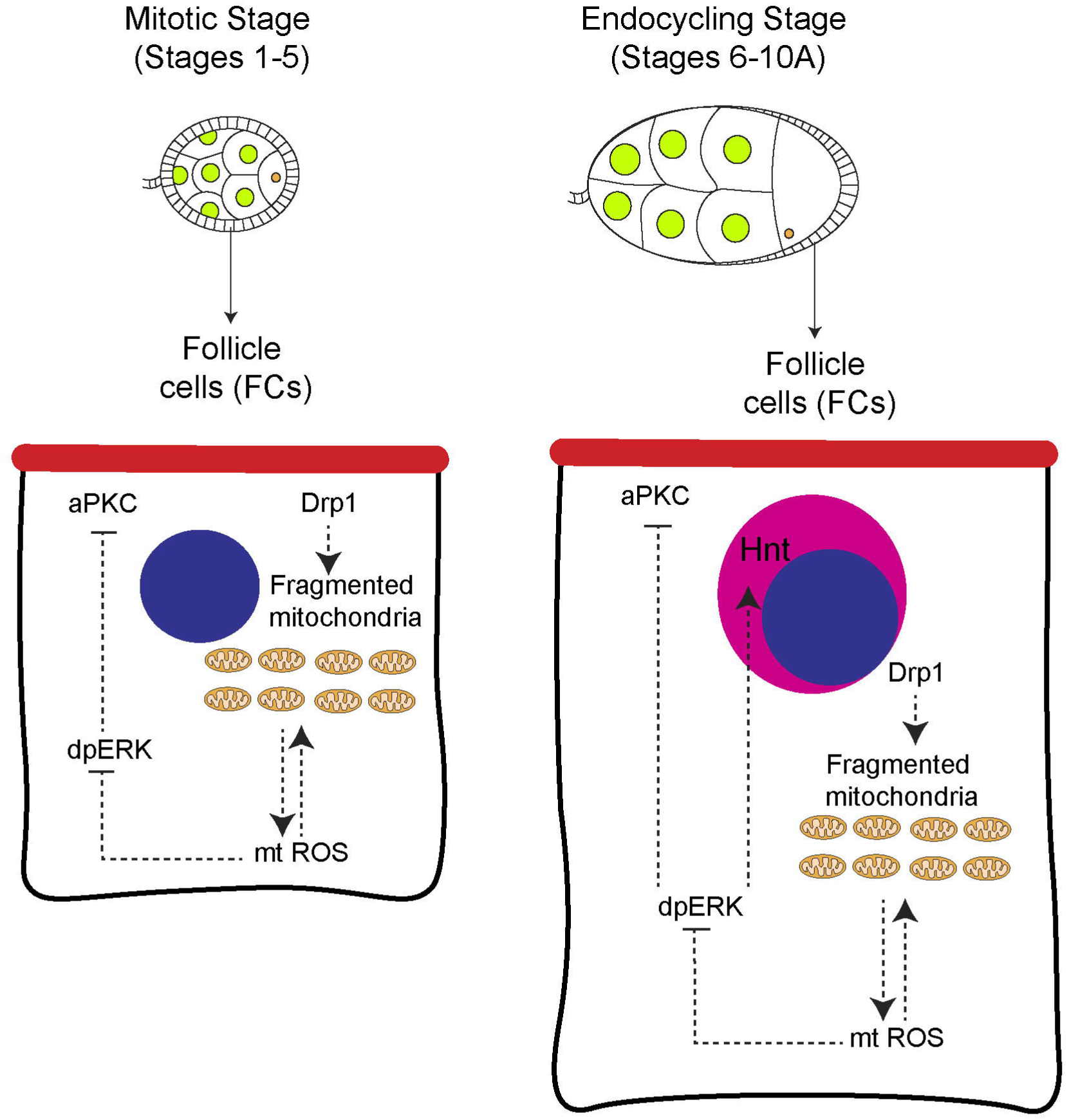
Interaction between mitochondrial morphology, ROS and EGFR and Notch signaling in *Drosophila* FC differentiation. The schematic shows the interaction between mitochondrial fission regulated by Drp1, ROS, and polarity protein aPKC in mitotic and endocycling FCs of *Drosophila* oogenesis. Drp1 regulates mitochondrial fission thereby controlling the levels of ROS and appropriate activation of dpERK in turn affecting the distribution of aPKC on the apical membrane in FCs in mitotic stages. In PFCs in endocycling stages, Drp1 leads to mitochondrial fission with ROS levels for appropriate activation of dpERK thereby regulating the accumulation of aPKC at the apical membrane and the activation of Notch signaling.

The apical polarity protein aPKC is decreased from the apical membrane in Drp1 depleted FCs in mitotic and endocycling stages. Appropriate recruitment of polarity proteins in stem cell proliferation is important for the correct orientation of the cell division machinery. aPKC regulates the mitotic spindle orientation and helps epithelial cell division in the *Drosophila* wing disc (Guilgur et al., 2012). However, aPKC is not required for correct spindle orientation in FCs (Bergstralh et al., 2013). However, aPKC depletion by optogenetic inactivation has been recently found to result in the formation of gaps and multilayered epithelia (Osswald et al., 2022). This occurs due to the increased activation of Myosin II. Loss of Myosin II activation in aPKC depleted FCs leads to an inhibition of actomyosin contractility (Osswald et al., 2022). It is therefore likely that an increase in actomyosin-based contractility occurs due to the decrease of apical aPKC in Drp1 depleted cells, leading to the occurrence of FCs in multiple layers. Future experiments testing the extent of increase in Myosin II activation in Drp1 depleted FCs will outline the mechanism by which multilayering occurs in the FC epithelium. A reverse process of loss of activation of Myosin II-driven contractility due to the decrease in ROS has been observed in Drp1 depleted *Drosophila* embryos during cellularization and dorsal closure (Chowdhary et al., 2020; Muliyil and Narasimha, 2014). Therefore, the effects on contractility due to mitochondrial fragmentation and ROS are likely to be tissue-specific and dependent upon the different factors that regulate Myosin II activity.

Alterations in mitochondrial ROS levels are known to occur on the change in mitochondrial morphology. An increase in mitochondrial ROS occurs on the depletion of Opa1 and a decrease occurs on the depletion of Drp1 (Celotto et al., 2012; Chowdhary et al., 2020; Quintana-Cabrera et al., 2021; Tang et al., 2009; Yarosh et al., 2008). In addition, an increase in ROS is also likely to impact mitochondrial morphology. For example, increase in ROS leads to mitochondrial fragmentation in cells undergoing delamination in dorsal closure in *Drosophila* embryogenesis (Muliyil and Narasimha, 2014). We also found that an increase in ROS due to the depletion of ROS scavenger proteins such as SOD2 and Catalase leads to mitochondrial fragmentation. ROS acts as a secondary messenger and can oxidize proteins and change their activity and distribution. An increase in ROS removes Opa1 from the mitochondria to the cytosol in mammalian H22T cells (Sanderson et al., 2015). This is likely to be a mechanism by which ROS elevation leads to mitochondrial fragmentation similar to Opa1 depletion. Decreased ROS during wound healing in epithelial cells leads to the inhibition of activation of Src kinase and inhibition of actomyosin constriction (Hunter et al., 2018; Muliyil and Narasimha, 2014; Ponte et al., 2020). ROS is known to regulate the activity of protein kinases and phosphatases (Corcoran and Cotter, 2013; Ray et al., 2012). ROS leads to oxidation of key catalytic cysteine residues and alteration of their enzymatic activity thereby affecting a variety of cellular processes. Therefore, an increase in ROS by the depletion of ROS scavengers in Drp1 depleted FCs is likely to oxidize and stabilize aPKC on the apical domain and correct epithelial polarity defects.

Our results showed that ROS increase in Drp1 depleted FCs not only led to the suppression of polarity defects but also decreased activated ERK and increased Notch signaling promoting the differentiation of PFCs. Multilayering of PFCs produced by depletion of polarity complexes often shows aberrant EGFR and reduced Notch signaling in the formation of PFCs. Depletion of the Dlg-Lgl-Scribble complex especially in the posterior leads to multilayering along with accumulation of dpERK and loss of Notch signaling (Li et al., 2008; Li et al., 2009; Tian and Deng, 2008). aPKC mutant clones show multilayering but their analysis for defects in EGFR and Notch signaling has not been reported in detail (Kim et al., 2009; Wodarz et al., 2000). Intestinal epithelial cells in *Drosophila* proliferate to form multilayers against bacterial infection and also show increased EGFR signaling (Buchon et al., 2010). EGFR and its downstream factors regulate the presence of fragmented mitochondria (Kashatus et al., 2015; Mitra et al., 2012). EGFR signaling is also known to require fragmented mitochondria produced by the activation of Drp1 driven fission for the proliferation of cancer cells (Kashatus et al., 2015). We found that increased ROS by depletion of Opa1 and ROS scavengers prevented the accumulation of dpERK. Decreased EGFR signaling and downstream factors are important for the formation of apical polarity in pre-FCs (Castanieto et al., 2014). This loss of dpERK is likely to be responsible for the stabilization of aPKC in the apical membrane in mitotic as well as endocycling stage FCs depleted of Drp1. It is further likely that aberrant EGFR signaling on the loss of polarity complexes also leads to a defect in mitochondrial morphology. Our data motivate a systematic analysis of the regulation of mitochondrial morphology proteins on the change in epithelial polarity complexes in FCs and other epithelial cells in *Drosophila*.

Increased EGFR signaling is known to oppose Notch signaling in various contexts such as the *Drosophila* wing disc and the FCs (Hasson et al., 2005). The transcription factor Groucho executes the repression between EGFR and Notch signaling in the wing disc. We have previously found that the Notch signaling-driven differentiation in FCs is promoted by the depolarization of mitochondria (Tomer et al., 2018). Moreover, a decrease in mitochondrial membrane potential also partially reverses the Notch-driven differentiation defect in Drp1 depleted PFCs (Tomer et al., 2018). We find that a decrease in ERK leads to restoration of aPKC and Notch signaling in Drp1 depleted PFCs. A link between apical polarity and Notch signaling has been shown in intestinal stem cells. Increase in aPKC in intestinal stem cells increases Notch signaling (Goulas et al., 2012). In this study, we find that mitochondrial fragmentation and increase in ROS driven by the depletion of Opa1 and ROS scavengers alleviates the Notch-driven differentiation defect in Drp1 depleted PFCs. In addition increase in ROS also leads to a rescue of the defect of accumulation of dpERK in Drp1 follicle cells. In summary, these studies together show that mitochondrial activity and dynamics along with appropriate ROS are important for mediating the antagonistic interaction between EGFR and Notch signaling in FCs.

## Acknowledgments

We thank Deepa Subramanyam, Mayurika Lahiri, Nagaraj Balasubramanian, and RR lab members for continuous discussions on the data. Stocks obtained from the Bloomington *Drosophila* Stock Center (NIH P40OD018537) were used in this study. We have used Microsoft Excel, Graph-Pad Prism, Fiji, and Image J for the analysis of data. We acknowledge the assistance provided by the *Drosophila* and Microscopy facilities for stock maintenance, media for experiments and microscopy. BU thanks the Council of Scientific and Industrial Research (CSIR), India for the graduate fellowship, and Science and Engineering Research Board (SERB), India for the project fellowship, and Innoplexus Consulting Services Pvt Ltd., and Scivic Engineering Pvt Ltd. for the additional financial support. RR thanks SERB, India for funding this project.

## Notes

### Competing Interest Statement

The authors have declared no competing interest.

